# High precision fluorescence tomography-guided system for pre-clinical radiation research: system design and validation

**DOI:** 10.1101/2025.07.03.662589

**Authors:** Xiangkun Xu, Yu-Pei Tseng, Hamid Deghhani, Luke Hardy, Zhishen Tong, Gao Guo, Amyn Habib, Iulian Iordachita, John W. Wong, Ken Kang-Hsin Wang

**Author notes:** **CORRESPONDENCE:** Xiangkun Xu, PhD, Department of Radiation Oncology, University of Texas Southwestern Medical Center, 2201, Inwood Road, Rm NC8.202, Dallas, TX, 75390, USA., Ken Kang-Hsin Wang, PhD, Department of Radiation Oncology, University of Texas Southwestern Medical Center, 2201, Inwood Road, Rm NC8.308E, Dallas, TX, 75390, USA.

## Abstract

**Purpose:** Cone-beam computed tomography (CBCT), commonly used for image guidance in pre-clinical studies, is limited in soft tissue localization and lacks the functional information needed for image-guided radiation studies and treatment assessment. To address these limitations, we developed 3D fluorescence tomography (FT) for high precision functional image-guided research and integrated it with small animal irradiators.

**Methods:** The FT system, integrated with a commercial bioluminescence-guided platform in a standalone configuration, enabling compact multi-projection and multi-spectral imaging. A dual-axis galvo mirror scanner was used for laser spots scanning to excite internal fluorophore. A transportable mouse bed allows animals imaged in the optical system and transferred to irradiators for CBCT imaging and FT-guided irradiation. Spatial geometry-based methods were developed to map the laser spots and fluorescence images onto the animal numerical mesh surface, generated from the CBCT image, to serve as input data for FT reconstruction. A self-calibration method using Born-ratio data, fluorescence normalized to the excitation image, was adopted for reconstruction. Mouse phantom embedded with QD800 reporter and orthotopic glioblastoma (GBM) model injected with IRDye 800CW labelled U87-EGFR cells were used to verify the FT-guided system in target localization.

**Results:** The laser spot mapping accuracy are within 0.7 mm maximum deviation. Our system can reconstruct the target QD800 as deep as 17 mm in the mouse phantom with single projection fluorescence imaging at localization accuracy < 1 mm deviation. Using the Born ratio approach, we effectively eliminated excitation light leakage and autofluorescence, enabling our FT system to localize IRDye 800CW-labeled EGFR-overexpressing GBM cells in the mouse brain with approximately 1 mm accuracy.

**Conclusion:** We developed a compact FT system integrated with a small animal irradiator, achieving sub-millimeter localization accuracy in phantom and *in vivo* models. Its precision, flexibility, and compatibility position it as a powerful tool for advancing functional imaging-guided pre-clinical radiation research.

## 1. INTRODUCTION

Preclinical study is an essential stage in radiation research, providing insights into new treatment strategies and radiobiological aspects before clinical trials^1^. Currently, cone-beam computed tomography (CBCT) is the primary imaging modality used for guiding radiation delivery on commercial small animal irradiators^2,3^. While CBCT provides valuable anatomical information, it is limited by low image contrast for soft tissue targets and lack of functional imaging capability, which can hinder precise tumor localization and accurate assessment of treatment outcomes. As such, there is a pressing need for more advanced imaging techniques to overcome these limitations in preclinical radiation research.

Optical imaging, including bioluminescence and fluorescence imaging, provides valuable insights into tumor biology and response to cancer treatment with high sensitivity and specificity at molecular and cellular level^4,5^. Besides, optical imaging systems are often more cost-effective and easier to operate compared to complex modalities such as magnetic resonance imaging (MRI) or positron emission tomography (PET). Our group and others have incorporated bioluminescence imaging with small animal irradiators to guide focal irradiation^6–9^. However, bioluminescence imaging was limited in providing functional imaging due to its reliance on specific luciferase-substrate systems. In contrast, fluorescent probes offer greater flexibility in terms of probe design and application, which makes fluorescence imaging become a powerful tool in visualizing biological process at the molecular level. Integrating fluorescence imaging with small animal irradiator would allow us to monitor the biological effect of radiation therapy, thus realizing biology-guided irradiation for preclinical research. However, widely-used two dimensional (2D) fluorescence imaging is inadequate in quantifying fluorophore distributions in vivo because of light diffusion in tissue. To overcome the limitation of 2D fluorescence imaging, three-dimensional (3D) fluorescence tomography (FT) was introduced to reconstruct the underlying fluorophore spatial distribution through a model of light propagation in conjunction of an optimization algorithm by minimizing the difference between calculated and measured surface fluorescence signal^10–14^. Recently, FT has been integrated with small animal irradiators, and the 3D localization and quantification performance were demonstrated with mice bearing 4T1 tumors^15^ and mice surgically implanted with ICG containing glass tube^16^, respectively.

To achieve highly accurate preclinical radiation study and facilitate the practical use of FT, we developed a compact FT system and integrated it a commercial bioluminescence-guided platform^8^ to significantly advance its imaging-guided capability. Compared to Shi et al.^15^, rotating animal in an upright position, and Nouizi et al.^16^, which rotates the imaging camera and fluorescence excitation laser in a C-arm irradiator, our system is a standalone unit featuring an innovative mouse bed system designed for compatible with irradiators for radiation guidance. The bed design allows the animal to lie naturally, preventing organ sagging compared to the upright set up^15^. Additionally, its standalone nature ensures seamless integration with existing irradiators, reducing optical alignment challenges^16^ while also enabling independent optical imaging applications. Moreover, to maximize the input data alleviating the ill-posedness of the inverse problem in FT, we utilized laser scanning to excite the imaged object and used single-projection detection in transmission imaging mode. This approach enabled us to generate multiple sets of fluorescence data for each laser scanning position. We further developed a novel method to map fluorescence images and laser spots on the animal surface to ensure accurate data input for the non-contact geometry of illumination and imaging setup. We also employed the normalized Born ratio^17^ for FT reconstruction by dividing fluorescence images by excitation images. This approach minimizes measurement uncertainties from excitation light leakage and autofluorescence, as well as reconstruction errors caused by optical heterogeneities in tissue, thereby improving target localization accuracy.

In this work, we demonstrated the localization performance of our FT system for deep seated target by embedding fluorescent probe QD800 to a mouse phantom. We further examined the functional imaging capability using an orthotopic glioblastoma model with over-expression of the epidermal growth factor receptor (EGFR). We expect that our FT-guided system will provide a powerful tool for pre-clinical radiation investigation.

## 2. MATERIALS AND METHODS

### 2.1 System configuration

Our FT system is integrated with the commercial bioluminescence-guided imaging platform MuriGlo (Xstrahl Inc., Suwanee, GA)^8^. As shown in Figure 1a, the FT module consists of a laser power control unit, a laser diode, a laser scanning control unit. The optical imaging chamber is included inside the MuriGlo. The power control unit includes a current controller (LDC 210C, Thorlabs, Newton, NJ) to adjust the laser output power and a temperature controller (TED 200C, Thorlabs) to stabilize the laser power by minimizing temperature fluctuation. The scanning control unit consists of a galvanometer (Galvo) system power supply (GPS011-US, Thorlabs), two driver cards (GVS002, Thorlabs), and a data acquisition device (NI USB-6211, National Instruments, Austin, TX). Inside the imaging chamber (Figure 1b and c), we integrated a fiber-coupled laser scanner, controlled by the Galvo system, into the MuriGlo for fluorescence excitation. The imaging chamber consists of a rotational 3-mirror system, a 4^th^ fixed 45° mirror (> 98% reflectivity), filter wheel (Edmund Optics Inc., Barrington, NJ), lens (50 mm, f/1.2, Nikkor, Nikon Inc., Melville, NY), and CCD camera (iKon-M934, Andor Technology, Belfast, UK). Fans (CFM-6015V-254-362-20, Digi-Key Electronics, Thief River Falls, MN) and a heater (FCH-FGC15132R, Omega, Norwalk, CT) are used to maintain the chamber temperature at desired level for experiments. The laser scanner (Figure 1c), including a dual-axis Galvo mirror (GVS002, Thorlabs) and a laser diode pigtailed with a single mode fiber (LP730-SF15, Thorlabs) and a collimator (PAF2-A4B, Thorlabs), was placed below the mouse bed. The scanner was utilized to steer the laser direction (pink arrows in Figure 1b) for fluorescence imaging. The laser diode-fiber configuration (Figure 1b and c) allows user easily changing the diode for desired fluorophore excitation without affecting optical alignment. The fluorescence signal emitted from the object surface was guided by the mirror system to the filter wheel and captured by the lens-CCD camera (white dashed line in Figure 1b). The image taken at top of the bed is labeled as 0° projection imaging. Our system allows laser excitation on the bottom of imaged object or animal through a transparent plate placing on the bed, while the fluorescence imaging can be detected from −90° to 90°projection relative to the imaged object. After optical imaging, users can detach the bed with the imaged object/animal and dock it to a small animal irradiator using a bed adaptor for CBCT imaging and 3D FT target reconstruction for radiation guidance^8^.

**Figure 1.**
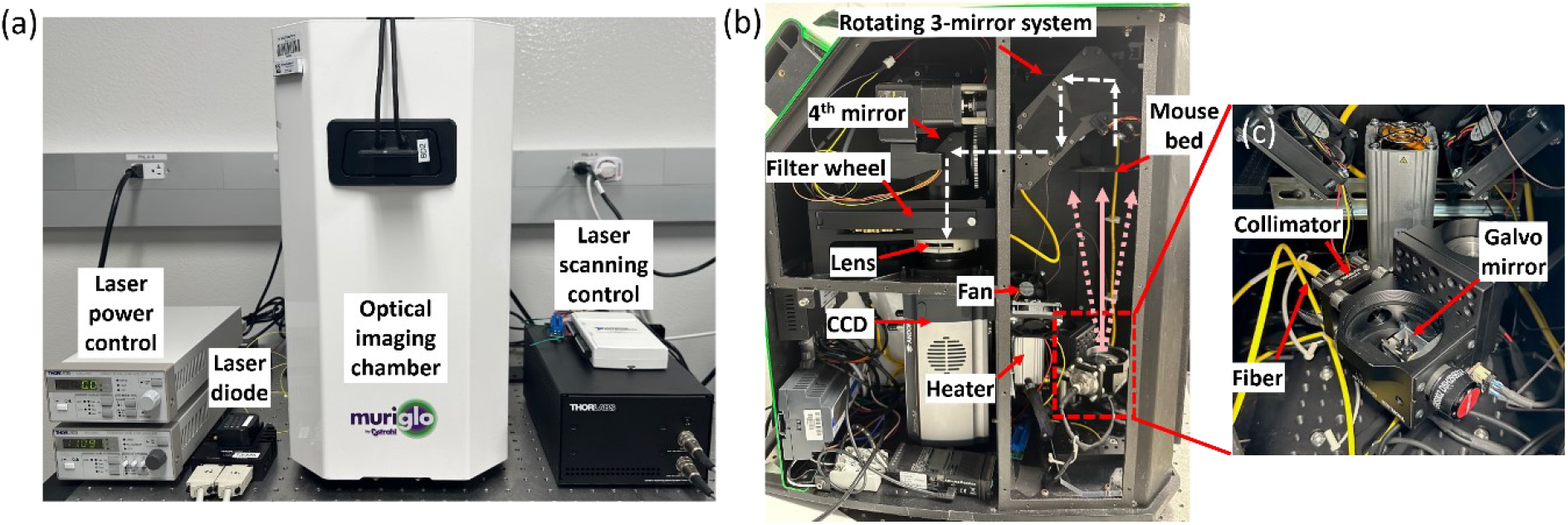
(a) shows the FT module including a laser power control unit, a laser diode, and a laser scanning control unit. The optical imaging chamber is part of the MuriGlo system. (b) illustrates the detail of the optical imaging chamber in (a). (c) is the laser scanner enlarged from the dashed area, depicted in (b).

### 2.2 Workflow of FT-guided system for irradiation

Figure 2 illustrates the workflow of using FT to guide irradiation in a small animal irradiator. First, surface fluorescence images are generated by positioning laser spot (LS) at a specific location on the imaged object to excite internal fluorophores. Photo images are then acquired at −60°, −30°, 0°, 30°, and 60° projections by rotating the 3-mirror system for geometrical calibration to register 2D CCD plane and 3D CBCT coordinate. After the optical imaging, the mouse bed along with imaged object will be transported to the irradiator, i.e. small animal radiation research platform (SARRP, Xstrahl Inc., Suwanee, GA) used in this work, for CBCT imaging. The CBCT image will be used to 1) provide anatomical structure of the imaged object to generate tetrahedral mesh for the finite element method (FEM)-based FT reconstruction and 2) define the 3D coordinate for treatment planning and irradiation. To implement FT reconstruction, we use an in-house developed automatic contouring method to generate binary object shape within region of interest, and the contour is used to generate the tetrahedral mesh with an averaged edge length of approximately 1 mm for FEM calculation. We then map the fluorescence/excitation images and laser spots to the mesh surface as the input for FT reconstruction. The excitation (Ex, Figure 2) image is acquired using excitation under the same geometric configuration as the fluorescence image. In conjunction with optical properties describing light absorption and scattering within the imaged object and reconstruction algorithm, we can retrieve the 3D spatial distribution of fluorophore for radiation guidance.

**Figure 2.**
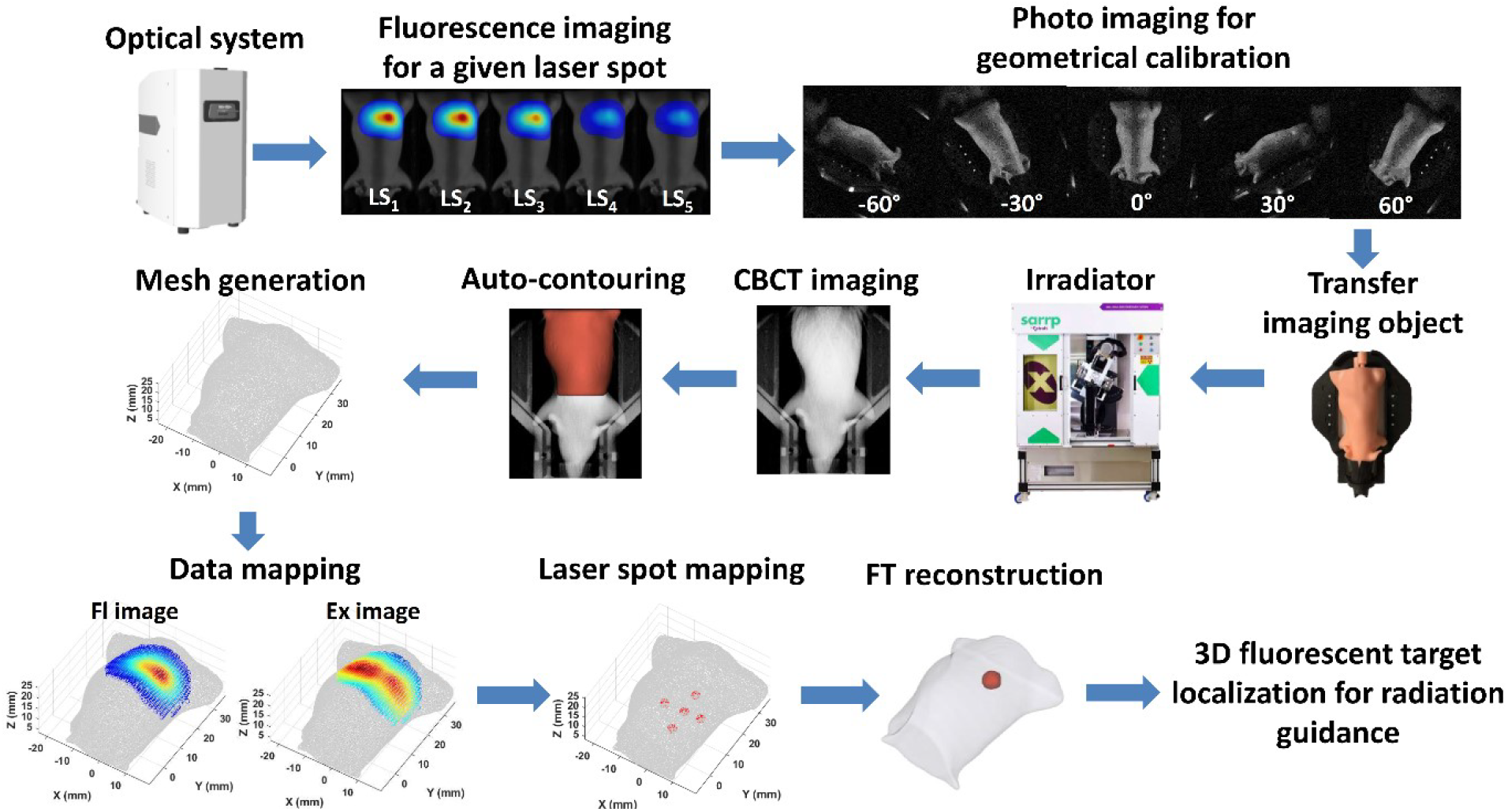
Workflow of fluorescence tomography-guided pre-clinical irradiation. The FI image refers to the fluorescence image. The excitation (Ex) image is acquired using laser excitation under the same geometric configuration as the fluorescence (FI) image.

### 2.3 Mapping fluorescence images and excitation images to object mesh surface

Because the CBCT image defines the coordinates used for FT reconstruction, we had proposed a geometric calibration method^8^ to register the 3D CBCT coordinate and 2D optical CCD plane to map the 2D optical images acquired at a given projection imaging angle onto the object mesh surface generated from the CBCT image. The details can be referred to our previous publication^8^. Briefly, in our geometrical calibration approach^8^, we firstly registered the 3D CBCT coordinate to 3D optical coordinates with rigid transformations, and then projected the 3D optical coordinate to 2D optical coordinates at the CCD plane using a pinhole camera model. These registration requiring the knowledge of geometrical parameters describing the relationship among different coordinate systems^8^. To retrieve these parameters, we used an optimization routine that minimizes the deviation between the measured and calculated positions of fiducial markers on the bed. The measured positions were obtained from 2D photo images (Figure 2), while the calculated positions on the photo images were derived using a calibration method^8^ and their coordinates in 3D CBCT space. After the 3D CBCT and 2D optical coordinates were registered, we can utilize the optimized geometrical parameters to map the optical signal from a given pixel of 2D fluorescence/excitation image to the corresponding position on the mesh surface of imaged object in 3D CBCT coordinate.

### 2.4 Mapping laser spots to object mesh surface

For precise FT reconstruction, the accurate laser spot position on the object surface is required. Because the transmission geometry was utilized for fluorescence imaging in our system, the laser beam was blocked by imaged object, which does not allow the CCD to capture the laser spot position on the object bottom surface. To determine the laser spot position, we utilized the intersection point between the laser beam and mouse bed (*LS_bed_*) as shown in Figure 3a1. The *LS_bed_* can be detected by the CCD when there is no object on the bed. With known *LS_bed_* and the spot position on the galvo system (*LG*), we can determine the laser beam, and then we can calculate the laser spot position on the object through the intersection between the laser beam and the object mesh surface.

**Figure 3.**
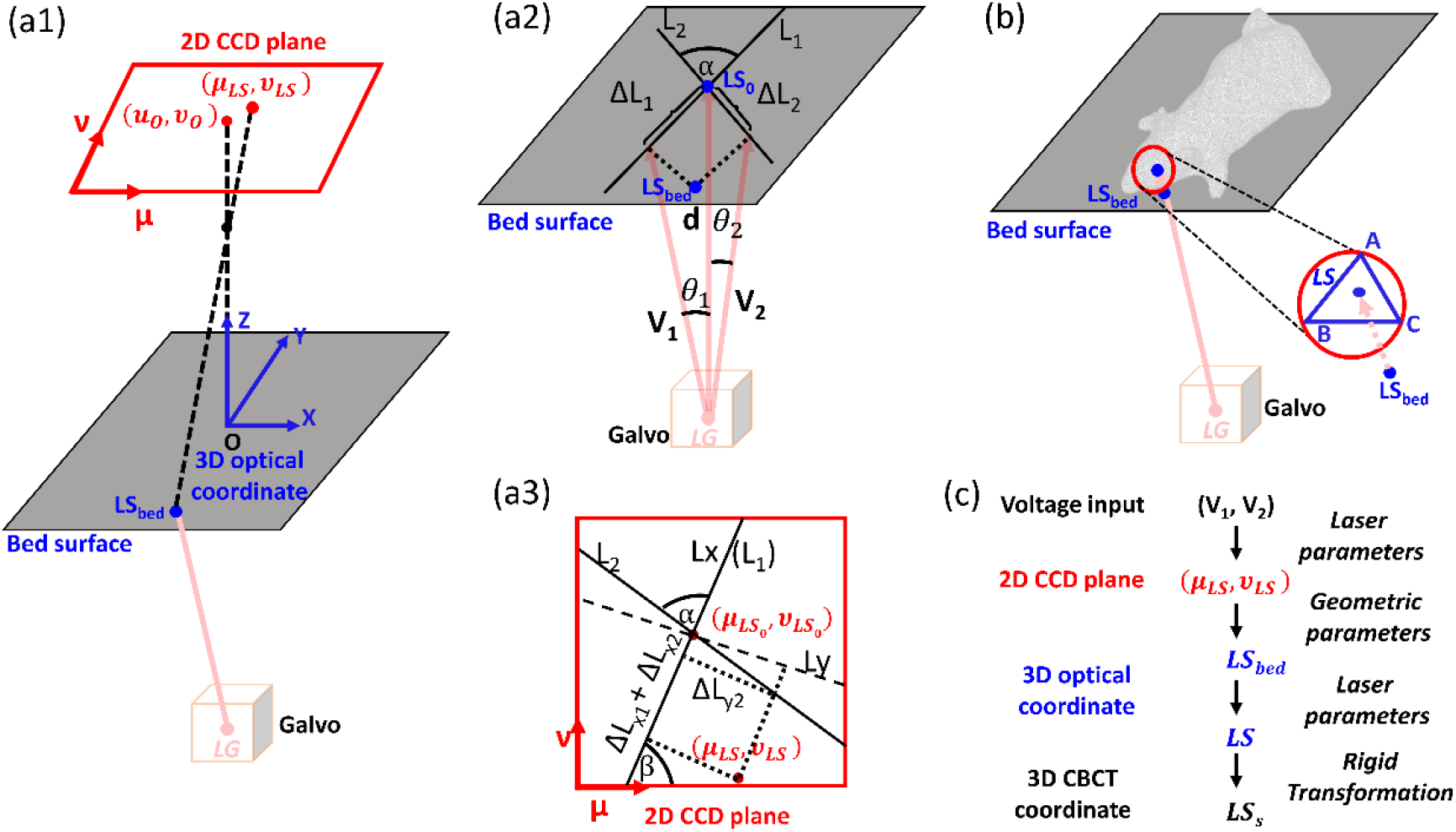
(a1) shows geometric relation among laser beam, galvo mirror, mouse bed and 2D CCD plane. The intersection *LS_bed_* between laser beam and mouse bed is projected to (*u_LS_, v_LS_*) on 2D CCD image plane through the pinhole camera model. (a2) shows the geometry between the dual-axis galvo mirror and mouse bed surface. When the laser beam is norm to mouse bed, the laser spot on the surface is defined as *LS_o_*. As the voltage *V_1_* or *V_2_* is applied, the laser beam will rotate *θ_1_* or *θ_2_* relate to 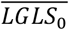 axis. We define *L_1_* and *L_2_* as the laser spot axis on the bed when *θ_2_* or *θ_1_* is 0 degree. The angle between *L_1_* and *L_2_* is *α*. (a3) show the geometric relation of the laser spots on the bed surface on 2D CCD plane. Two orthogonal *L_x_* and *L_y_* axes are used for the orthogonal decomposition of the distance shift of laser spot relative to 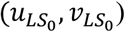. (b) shows the geometry relation of *LG, LS_bed_*, and the laser spot position *LS* on the mesh surface. (c) shows the overall workflow of mapping laser spots to object mesh surface in 3D CBCT coordinate from voltage input on the galvo mirror.

The *LS_bed_* depends on the input voltages of the galvo mirror and the geometry parameters of laser system. To determine *LS_bed_*, we model the geometry of laser beam intersecting with the mouse bed surface (Figure 3a2-a3). For a given voltage (*V*_1_, *V*_2_) of the dual-axis galvo mirror, the mirror rotation angle can be written as:

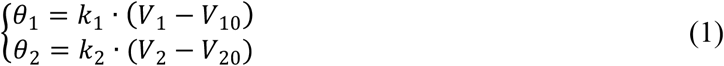

where *θ*_1_ and *θ*_2_ are the angle between laser beam and 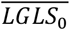 axis. 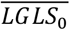 axis is when the laser beam is normal to the bed with input galvo voltage as (*V*_10_, *V*_20_). The distance between *LG* and *LS_o_* is d. The parameter *k_1_* and *k_2_* are the constant factors transferring from galvo voltage to the angle *θ*. The distance of the laser spots between *LS_o_* and *LS_bed_* along the line *L_1_* and *L_2_* can be written as:

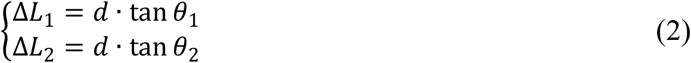

where *L_1_* or *L_2_* is the laser trajectory on the mouse bed when the input *V_1_* or *V_2_* is changing and *V_2_* or *V_1_* is constant as *V_20_ or V_10_*, respectively. *LS_o_* and *LS_bed_* can be expressed as 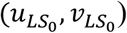 and (*u_LS_, v_LS_*) on the 2D CCD plane (Figure 3a3), respectively. We then introduced Cartesian coordinate system *Lxy* in the CCD plane where *Lx* axis is overlapped with *L_1_* to calculate the position of laser spots *LS* in (*u, v*) coordinate. In the *Lxy* system, the coordinate of laser spot *LS* can be expressed as:

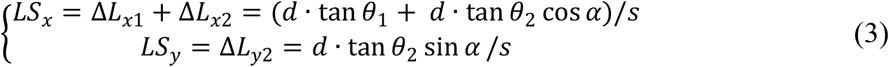

where *s* is the pixel scale, 0.102 mm/pixel for MuriGlo. We then translate *LS* position from the *Lxy* coordinate to the CCD coordinate with angle *β*,

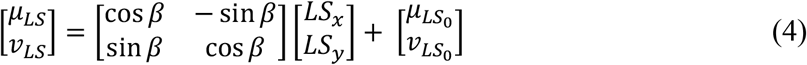

For a given voltage (*V*_1_, *V*_2_), we can calculate the laser spot position (*u_LS_, v_LS_*) on the 2D CCD image plane with the 9 parameters 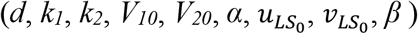 and Eqs. (1-4).

To visualize laser spots, we placed a diffuse plate on the mouse bed. To obtain these laser geometric parameters, the diffuse plate was placed at two different heights (0 mm and 11.4 mm) above the bed surface, and the positions of laser spots on the plate were measured through CCD imaging when different input voltages were applied to the galvo mirror. The positions of laser spots can also be calculated based on the input voltage and the Eqs. (1-4). An optimization program with a constrained multivariable optimization function (fmincon; MATLAB R2019b, The MathWork Inc., Natick, MA) was developed to retrieve the laser parameters by minimizing the deviation between the calculated and measured laser spots on the CCD image plane.

With the parameters in place, we can determine the laser spot position on the object surface for any given voltage (*V*_1_, *V*_2_) applied to the galvo mirror. The position of *LS_bed_* in 3D optical coordinate was determined through mapping its projected position (*u_LS_, v_LS_*) in 2D CCD image to the bed surface with the pinhole camera modelling used in the geometrical calibration. In 3D optical coordinate *O-XYZ*, the coordinate of (*u_LS_, v_LS_*) shown in Figure 3a1 can be expressed as:

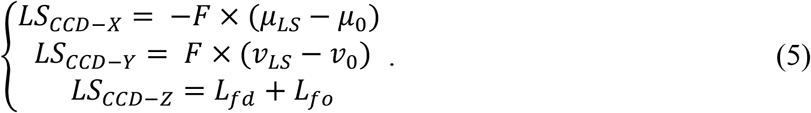

where *L_fd_* is distance from the focal point, *f*, to the 2D CCD plane; *L_fo_* is distance from the focal point, *f*, to the 3D optical coordinate origin *O*; *F* is physical pixel size, 13 μm/pixel, of the CCD sensor array; and (μ_0_, ν_0_) is center of the 2D optical image, intersection between the axis *Z* in *O-XYZ* with CCD image plane. These parameters (μ_0_, ν_0,_ *L*_*fd,*_*L*_*fo*)_were determined through the geometrical calibration process as described in Section 2.3. The coordinate of focal point *f* can be written as (0, 0, *L_fo_*) in the 3D optical coordinate *O-XYZ*. Thus (*u_LS_, v_LS_*) and *f* can form a ray. We can use the Möller-Trumbore ray-triangle intersection algorithm^18^ to locate the intersection *LS_bed_* within the bed surface. With the same method, we can also locate *LS_o_* on the mouse bed when the laser beam is normal to the mouse bed. Therefore, we can determine the *LG* in 3D optical coordinate *O-XYZ* based on the distance d between LG and mouse bed.

With known *LG* and *LS_bed,_* we employed the Möller-Trumbore ray-triangle intersection algorithm again to calculate the laser spot position (*LS*) on the mesh surface, the intersection between the laser ray formed by *LG* and *LS_bed_* and a surface triangle ABC of an imaged object mesh, as shown in Figure 3b. The barycentric coordinates is utilized in the Möller-Trumbore algorithm, which we can write the coordinate of the *LS* as

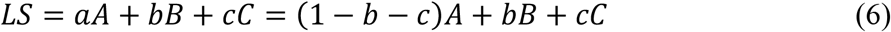

*A, B* and *C* are the coordinates of the vertices for a triangle *ABC* in 3D optical coordinate *O-XYZ* (Figure 3b) and *a, b, c* are the barycentric coordinates of *LS* for the triangle *ABC.* The intersection *LS* can also be represented by the ray from *LS_bed_* to *LS* as

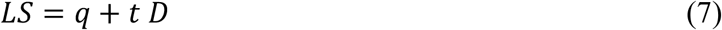

where *t* is the distance from *LS* to *LS_bed_*, and *D* is the normalized direction of the ray connecting *LG* and *LS_bed_*. By combining Eqs. (6) and (7), the unknowns *t, b* and *c* can be solved.

With the above-mentioned method, we can identify the barycentric coordinates, *a, b*, and *c* of the *LS* for the triangle ABC in the 3D optical coordinate. Since the 3D CBCT and optical coordinates are registered through rigid transformation, we can apply the barycentric coordinates, *a, b*, and *c* to the corresponding triangle *AsBsCs* in the 3D CBCT coordinate with

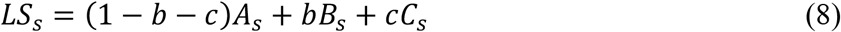

For a given closed 3D mesh surface, there will be more than one intersection. The point *LS_s_* we chosen is the closest intersection to *LS_bed_*.

To evaluate the accuracy of the laser spot mapping to CBCT coordinate, we designed a diffuse step phantom. Fiducial markers in the form of 1.5 mm diameter holes were incorporated into the phantom to allow us locate the laser spots in both optical and CBCT images. The step has a height of 4 mm, and the thickness of the step is 1.1 mm, consistent with the thickness of the transparent plate supporting the animal. The dimension of the phantom is same as the supporting transparent plate, allowing it to be easily placed on the bed. The laser spots on the mesh surface of the phantom in CBCT coordinate can be calculated through the workflow in Figure 3c, based on the input voltage and the optimized laser parameters. The measured laser spots in CBCT coordinate were determined by the relative position of laser spot and its surrounding fiducial holes in 2D CCD image plane and the coordinates of these fiducial holes in CBCT image. The accuracy of laser spot mapping is evaluated by the deviation between the calculated and measured laser spots in CBCT coordinate.

### 2.5 Mathematical framework for FT reconstruction

Because light transport in tissue is dominated by multiple scattering, two diffusion equations were applied to model the fluorescence at emission wavelength (*λ_m_*) generated by a fluorescent agent excited by an external light source at excitation wavelength (*λ_x_*):

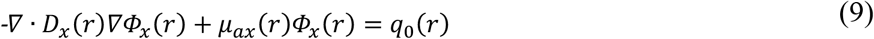

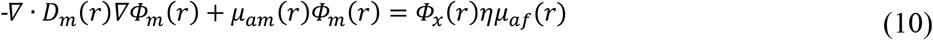

where Φ_*x,m*_(*r*) is the excitation (*x*) or fluorescent (*m*) photon fluence rate at location *r*, and 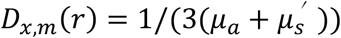 is the corresponding diffusion coefficient at location *r* at wavelength *λ_x_* and *λ_m_*, respectively. *μ_a_* and 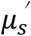 are absorption and reduced scattering coefficients. *q*_0_ (*r*) is the excitation source distribution. *ημ_af_*(*r*) is the fluorescence yield, a product of the quantum efficiency *η* and the absorption coefficient *μ_af_* of fluorescent agent. An index-mismatched type III condition was used to describe the air-tissue boundary.

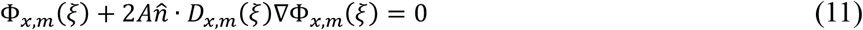

*ξ* represents points on the tissue boundary, and coefficient *A* can be derived from Fresnel’s law, depending on the refractive index of tissue and air. 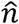 is the unit vector pointing outward, normal to the boundary.

The goal of FT reconstruction is to recover the unknown fluorescence yield *ημ_af_* by minimizing the difference between measured and calculated fluorescence data from animal surface. The FT reconstruction is an iterative optimization process, leading to an update equation given by^19^:

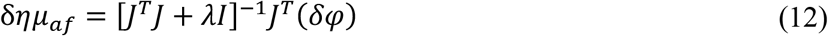

where *δφ* is the difference between the logarithm of measured surface fluence rate 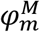 and calculated surface fluence rate 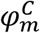 at each iteration, *λ* is the regularization parameter, and *I* is an identity matrix. The Jacobian matrix, *J*, is the sensitivity matrix that relates a small change in the logarithm of boundary fluence rate to a small change in fluorescence yield *ημ_af_*, and it can be written as^20^:

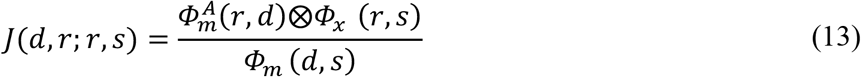

Where, 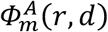 is the adjoint emission field at position *r* from adjoint source at a detector position *d*; *Φ_x_* (*r, s*) is the excitation field at position *r* from an excitation source position *s*; and *Φ_m_* (*d, s*) is the emission field at detector position *d* from an excitation source position *s*. The Jacobian matrix is generated by NIRFAST package^21^ under finite element framework, and the convolution ⨂ is used to account for the effect of non-homogeneous field distribution for each spatial node and its associated neighboring nodes. In the case of Born ratio data, where measured emission surface fluence rate 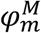 is scaled by the measured excitation surface fluence rate 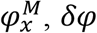 can be written as 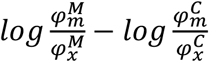.

### 2.6 Mouse phantom experiment

To access FT localization accuracy, we utilized a fluorescent mouse phantom (XFM-2, Perkin Elmer Inc., Waltham, MA) made of polyurethane material that includes scattering particles and dye to simulate optical properties of live tissue. A rod containing a fluorescent probe QD800 embedded in the tip was inserted into the phantom at the depth of 7 mm and 17 mm separately to simulate an internal fluorescent source at various depths.

The laser diode at 633 nm (LP633-SF50, Thorlabs Inc) was used for excitation, and 5 laser spots with diameter less than 1 mm and spatial relative distance around 5-7 mm were sent individually onto the bottom of the phantom for fluorescence imaging. The excitation and fluorescence images were acquired from the phantom top surface when the 3-mirror system is at 0° with an excitation filter at 630 nm (20 nm bandwidth, ET630/20x, Chroma Technology Corp., Bellows Falls, VT) and an emission filter at 800 nm (40 nm bandwidth, AT800/40m, Chroma Technology Corp., Bellows Falls, VT), respectively. Because of the strong excitation signal, 1 × 1 binning and exposure time < 1 second was utilized to not saturate the CCD detector for excitation images. Binning size at 8 × 8 and exposure time of 1-2 seconds was utilized for fluorescence images. After the fluorescence imaging session, photo images at −60°, −30°, 0°, 30° and 60° projections were acquired to obtain the geometric parameters for mapping fluorescence or excitation images to object mesh surface. The camera setting of 4X pre-amplifier gain and 1 MHz readout rate was used for all the optical data acquisition.

After optical imaging, the mouse phantom with the detachable mouse bed was docked to the bed adaptor pre-installed in SARRP for CBCT imaging. A region of interest was cropped from the CBCT image and used to create a contour for mesh generation. We used NIRFAST package^21^ and CBCT image to generate tetrahedral mesh with an average edge length of 1 mm for FT reconstruction. Fluorescence/excitation images and laser spots were mapped to the mesh surface through the data mapping method described in Section 2.3 and 2.4, respectively. After data mapping, a threshold of 10% of the maximum mapped fluorescence data value was used to select the input data for FT reconstruction to remove noisy low value data. We investigated the FT localization accuracy using both fluorescence data only and Born ratio method by comparing to the actual fluorescence probe location, where can be determined from CBCT image. The FT reconstructed source was delineated by a threshold of 0.5 of maximum reconstructed value. The localization of reconstructed source was determined by the center of mass (CoM) of FT reconstructed source. The values of *μ_a_* 0.005 and 0.008 mm^−1^ and 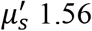 and 1.87 mm^−1^ for 633 and 800 nm, respectively, and refractive index 1.5, provided by Perkin Elmer, were used for the reconstruction.

### 2.7 *In vivo* experiment with GBM model

*In vivo* procedures were carried out in accordance with the institutional animal care and use committee at the University of Texas Southwestern Medical Center. To demonstrate the localization capability of our FT system *in vivo*, we established GBM model with epidermal growth factor receptor (EGFR) over-expressed brain tumor cells U87-EGFR^22^. The U87-EGFR cells were labelled with IRDye 800CW EGF optical probe (LI-COR, Lincoln, Nebraska) by mixing the cells at 1 × 10^5^ cells/μL with the probe at 10 *p*mol/µL in phosphate buffer solution at volume ratio of 50:1. We implanted 8 × 10^5^ IRDye labelled U87-EGFR cells into the left striatum of nude mice (J:NU 007850, The Jackson Laboratory, Bar Harbor, Maine) at 3 mm depth from a surgical opening.

*In vivo* fluorescence imaging was conducted 48 hours after cell implantation. Considering the excitation and emission spectrum of IRDye 800CW, laser diode at 730 nm was used for excitation, and fluorescence signal was collected with filter at 800 nm and 40 nm bandwidth. Laser spots were sequentially placed to the inferior side of mouse head with relative spacing around 3.5 mm, and 7 spots in total were used. Fluorescence images and excitation images were acquired from the top surface of mouse head when the 3-mirror system was set at 0° angle. Because of the strong transmission signal from excitation laser, besides the camera setting of 1×1 binning and < 1 second exposure time, we further used OD2 filter (NDUV2R20A, Thorlabs Inc) to attenuate the excitation signal to avoid image saturation. The camera setting of 4X pre-amplifier gain and 1 MHz readout rate was used for data acquisition. Photo images at −60°, −30°, 0°, 30°, and 60° projections were acquired for geometric calibration. The imaged mouse was anesthetized with 2% isoflurane in oxygen through the nose cone, and the rubber bands and taps on the bed were used to secure the limbs and tails. After the optical imaging, we transferred the mouse with the detachable bed to the bed adaptor pre-installed in SARRP for CBCT imaging. Due to excitation light leakage into the fluorescence detection window and auto-fluorescence from the mouse brain, using a 10% threshold of the maximum fluorescence signal to select input data for FT reconstruction proved insufficient. Thereafter, the value of Born ratio data larger than an animal-specific threshold was used to remove the data points where excitation leakage or auto-fluorescence dominated for FT reconstruction. A threshold of 0.5 of maximum FT reconstructed value was used to delineate reconstructed target. We used *μ_a_* 0.025 and 0.032 mm^−1^ and *μ_s_’* 1.29 and 1.17 mm^−1^ at 730 nm and 800 nm, respectively, and refractive index 1.4 for FT reconstruction.

To validate the accuracy of FT localization, T_2_-weighted fast spin echo sequence magnetic resonance imaging (MRI) (MRS 3017, 3T, MR Solutions Ltd. GU3 1LR, UK) was utilized to define the cell injection position of GBM-bearing mice as the ground truth. The MRI and SARRP CBCT image of the mouse head were manually registered using 3D slicer (version 4.11.2; https://www.slicer.org). The deviation between CoM of FT reconstructed IRDye distribution and the cell injection position was utilized to evaluate FT localization accuracy.

## 3. RESULTS

### 3.1 Validation of laser spots mapping

The procedure and validation of laser spot mapping to object surface is demonstrated in Figure 4. Figure 4a shows the results of laser parameters optimization through measuring the laser spots on the diffuse plate in the 2D CCD coordinate when the plate was placed at 0 and 11.4 mm above the mouse bed surface corresponding to 8 pairs of input voltage on the galvo mirror. The laser parameters were optimized through minimizing the deviation between measured and calculated positions of laser spots. The average and maximum deviation among all laser spots after optimization is at 0.12 mm and 0.28 mm, respectively. Figure 4b shows the validation of mapping laser spots to the step phantom surface in 3D CBCT coordinate. The mapped position was calculated from the procedure in Section 2.4 with the input voltage and the optimized laser and geometric parameters. The reference positions were determined from the measured laser spots on the step phantom surface. The average and maximum deviation between the mapped and reference positions of laser spots on the step phantom surface are at 0.32 and 0.62 mm, respectively. This result indicates we can map laser spots to object surface in 3D CBCT coordinate for a given input voltage at sub-millimeter accuracy.

**Figure 4.**
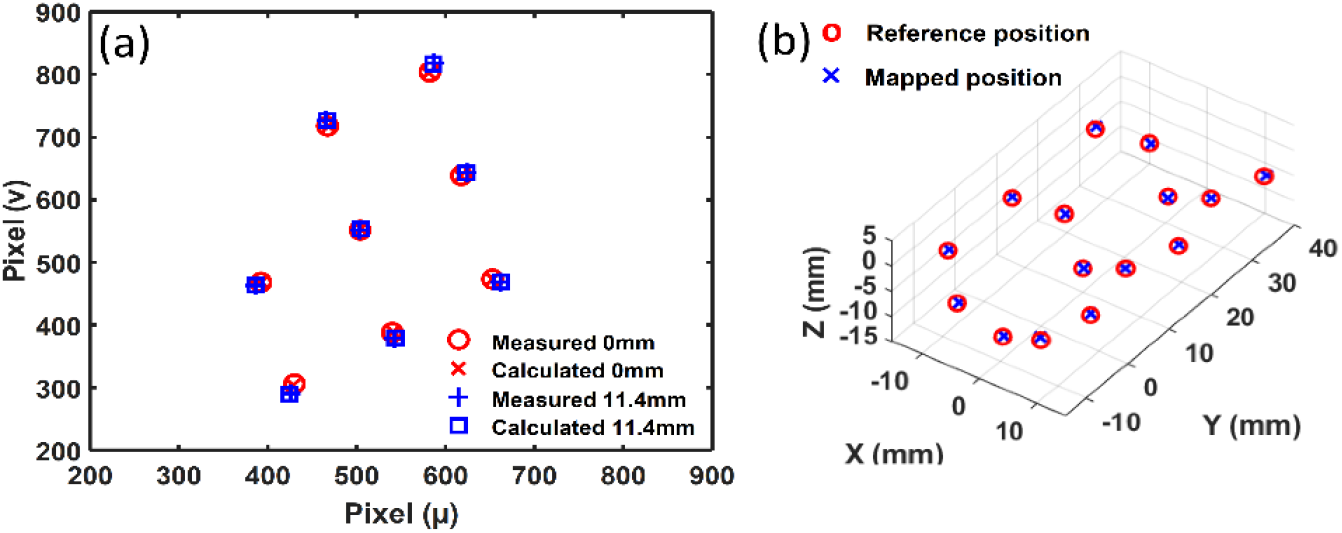
Validation of laser spots mapping; (a) shows the results of laser parameter calibration by placing the diffuse plate at 0 and 11.4 mm above the mouse bed. (b) Validation of laser spot mapping to 3D CBCT coordinate; red circles represent the measured laser spots positions on the step phantom surface in 3D CBCT coordinate, and blue crosses represent the mapped positions of laser spots on the step phantom surface.

### 3.2 Fluorescence tomographic image of deep-seated fluorophore QD800 in mouse phantom

To evaluate the target localization accuracy of our FT system at different depths, we used a tissue-mimicking mouse phantom embedded with the fluorophore QD800. A single fluorophore probe was positioned at depths of 7 mm and 17 mm in the phantom for two independent fluorescence measurements, as illustrated in Figure 5a. Figure 5b1 and b2 are the representative fluorescence image acquired at 0° projection mapped to the phantom and its mesh surface, respectively, when the fluorescent target was placed at depth of 7 mm and the corresponding laser spot set on the bottom of the phantom is shown in Figure 5b3 (pointed out by the blue triangle). Total 5 laser spots for a given target depth (red circles, Figure 5b3) led to 5 fluorescence images used for image reconstruction. The representative fluorescence images and laser spot for the target at depth of 17 mm are shown in Figures 5c1-c3.

**Figure 5.**
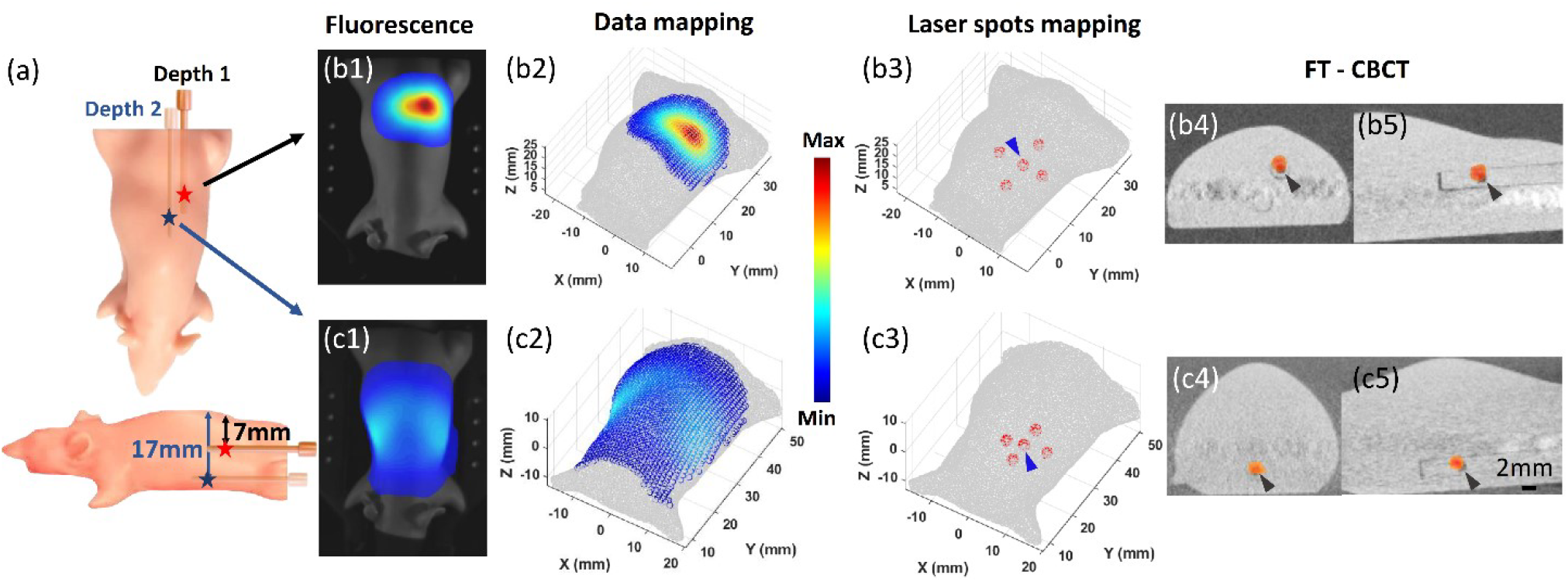
Phantom FT results; (a) shows the fluorophore QD800 (indicated by asterisk) buried in a rod was inserted into the mouse phantom at depth of 7 mm and 17 mm for two independent measurements. (b1-b5) and (c1-c5) show the fluorescence image at 0° projection, the image mapped to mesh surface, laser spots mapping and FT reconstruction results (heat map) overlapped with CBCT image for the scenarios where the fluorophore was at depth of 7 and 17 mm, respectively. The blue arrow in (b3) and (c3) indicated the laser spot position for fluorescence image (b1) and (c1), respectively. Data > 10% of maximum mapped valued is displayed. The black arrows in (b4-b5) and (c4-c5) indicated the actual position of fluorophore QD800.

We compared the CoM of FT delineated target to the ground truth position of the fluorophore pointed by the black arrow shown in CBCT image (Figure 5 b4-b5 and c4-c5), the deviation was at 1.0 and 0.9 mm when the fluorophore was at depth of 7 and 17 mm, respectively. The results demonstrated our FT system can reach the localization accuracy at 1 mm even for deep-seated targets. We further investigated the FT localization using Born ratio data, the deviation between the CoM of the FT reconstructed target and the fluorophore ground truth was at 1.2 and 1.5 mm (results not shown) when the fluorophore was at depth of 7 and 17 mm, respectively.

### 3.3 FT reconstruction of *in vivo* EGFR over-expressed GBM

We used a GBM model with EGFT over-expressed cells to assess the FT reconstruction performance *in vivo*. Specifically, the effectiveness of Born ratio in aiding *in vivo* FT accuracy is clearly illustrated in this case. We observed the fluorescence images (Figure 6 a1 and b1) contaminated from excitation signal (Figure 6 a2 and b2) because of the excited laser leaked into the emission window. This is especially evident when the laser spots were placed at thinner part of animal body, such as close to sides of the brain (Figure 6 b1 and b2 resulted from laser spot 2, LS_2_, Figure 6c). The light contamination from excitation thus confounds the fluorescence signal (Figure 6 a1 vs a2 and b1 vs b2). By applying Born ratio imaging, the ratio of fluorescence and excitation image (Figure 6 a3 and b3), we can suppress the contribution of leaked excitation signal. Figure 6 a3 and b3 show the improved contrast of fluorescence signal in the Born ratio images, which shows the image contrast highly depend on the excited laser spot location. To improve the FT reconstruction accuracy, we used a threshold of 0.01 for ratio images to remove the data points contaminated from excitation light and autofluorescence. The chosen data used for FT reconstruction were shown in Figure 6 a4 and b4 for the cases of laser spot 1 and 2, respectively, shown in Fig. 6c. By comparing Figure 6 a4 and b4, we can see the pattern of the Born ratio images changed with the laser spot location. The fluorescence tomographic image was overlapped with MRI image to validate the localization accuracy (Figure 6 d-e). Because of the small amount of injected cells, it was challenging for the EGFR over-expressed tumor cells to be identified in the MRI. Instead, we used the tumor injection site as the ground truth to validate the FT localization accuracy. The localization accuracy of our FT system is at 1.1 ± 0.4 mm (n = 4) by comparing the CoM of FT reconstructed target and the tumor injection site.

**Figure 6.**
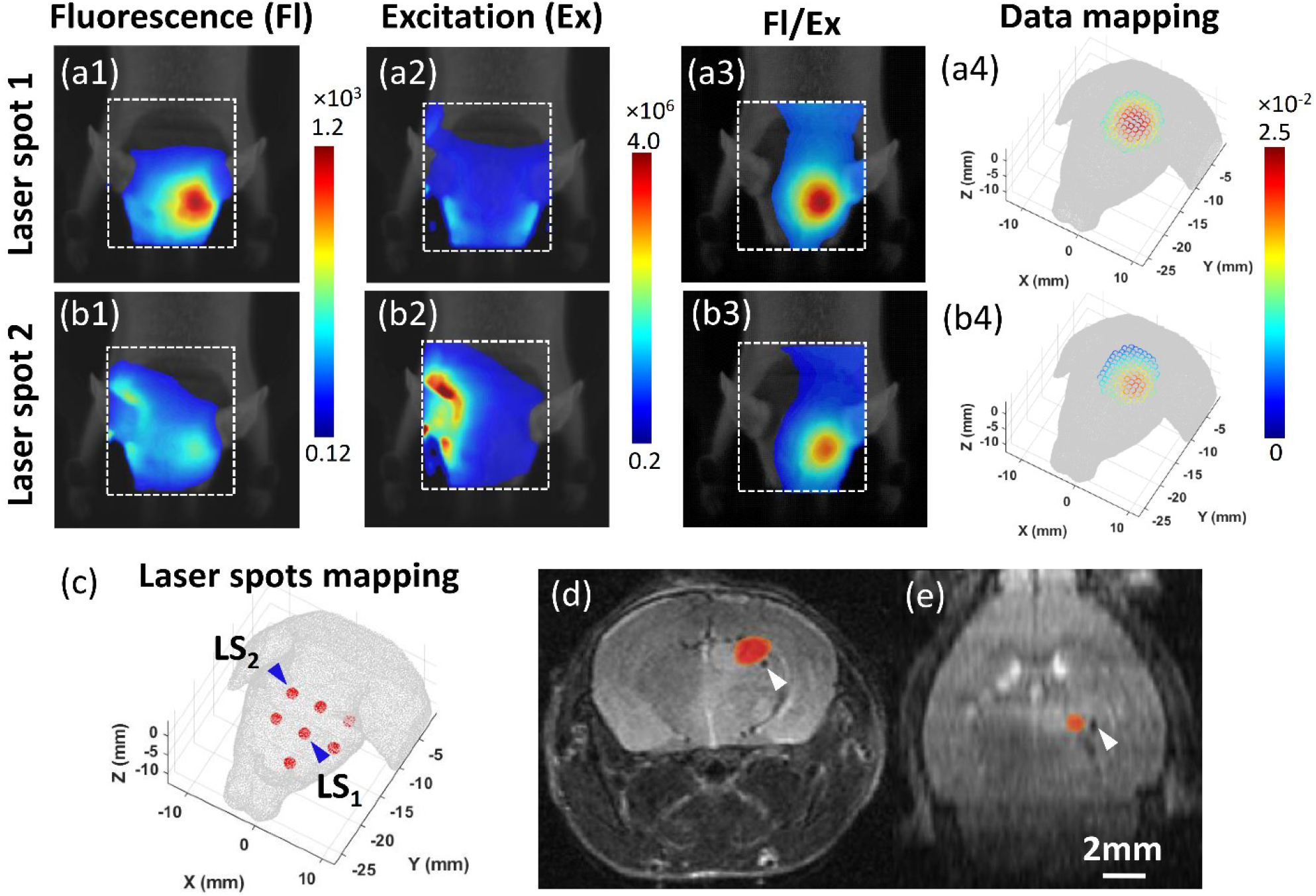
*In vivo* results; (a1-a3) and (b1-b3) shows the representative images of fluorescence, excitation, and ratio between fluorescence and excitation signal for laser spot 1 and laser spot 2, respectively. The white dashed region indicates the region of interest for display. (a4) and (b4) show the data mapping of ratio images on mesh surface from (a3) and (b3), respectively. Only data used for fluorescence tomography reconstruction was shown. (c) shows the seven laser spots used for fluorescence imaging were mapped to the mesh bottom surface. LS_1_ and LS_2_ pointed out the position of laser spot 1 and laser spot 2, respectively. (d-e) are transverse and coronal views of MRI image overlapped with fluorescence tomographic image. The white triangle points out the cell implantation site. A threshold of 0.5 was applied to delineate the fluorescence reconstructed target (heat map).

## 4. DISCUSSION

It is worthwhile to emphasize that the integration of FT with small animal irradiators brings two major advantages to preclinical radiation therapy: precise target localization and access to functional information. The level of precision in pinpointing the target is critical for delivering radiation therapy with high accuracy, especially in soft tissue environment where commonly used CBCT on small animal irradiators falls short. Small animal radiation research would also be greatly enhanced with capabilities of biologic and functional targeting and assessment beyond anatomical imaging. Fluorescence imaging and tomography would allow in-depth biological and mechanistic interrogation of key information about the tumor and its microenvironment such as hypoxic burden and metabolic rate, etc., that are known to impact response to RT. It complements our previously developed bioluminescence imaging^8,23^ to study tumor and normal tissue response, when the use of engineered bioluminescent models is not available. Fluorescence imaging also has an added advantage to be amenable to clinical translation as a surrogate for PET imaging^24,25^. The research potentials of FT are particularly significant when considering combinational molecular therapeutic strategies with radiation, such as immunotherapy. Integrated CBCT/FT capabilities that complements the small animal irradiator would provide a powerful platform for pre-clinical radiation research.

In our FT system, we utilized transmission imaging geometry by allowing laser excitation from the bottom surface of the imaged object and collecting fluorescence signal from the top surface. With animals placed in a nature comfort position on the bed for optical imaging, it eases the requirement for animal immobilization and transport between the optical system and irradiators. Our previous study^26^ shows the positioning error can be maintained within 0.2 mm as long as the animal was effectively immobilized and anesthetized during transportation. Compared to the systems that rotating animals^15^ or cameras^9,16^, our design simplifies the imaging process by avoiding the challenges of upright animal positioning and reducing mechanical complexity. Although single-projection imaging geometry is sufficient for the phantom (Figure 5) and *in vivo* studies (Figure 6) shown in this work, our system can support multiple projection imaging by rotating the 3-mirror assembly (Figure 1b) which is significant simpler than rotating entire camera^19^ when multiple views are required. The laser scanning system (Figure 1c) also increases data input by enabling different fluorescence surface patterns for a given laser excitation spot (Figure 6 a1, b1 vs. c), helping to mitigate the ill-posedness of the inverse FT problem. Furthermore, compared to other systems^15,16^, the standalone feature of the optical system with the transportable bed minimizes the need of modifying existing irradiators and ensures fluorescence imaging-only applications without occupying the irradiator that affects its experiment throughputs.

With the integration of FT into the commercial platform MuriGlo, we can utilize the previous published method for mapping optical images to the object mesh surface with the mapping error within 0.7 mm^8^. We further innovated the laser spots mapping method (Figure 3). We calibrated the laser scanning parameters through a diffused plate to effectively capture the laser spots using the camera. With the laser scanning parameters and data mapping geometric parameters in place (Section 2.4), we can accurately calculate the laser spot positions on the object mesh surface within 0.7 mm accuracy (Figure 4b), which is important to maintain high precision target localization of the FT-guidance for irradiation.

Based on the accurate data and laser spot mapping methods, and the diffusion approximation reconstruction approach (Section 2.5), we evaluated the FT system performance using both a mouse phantom and *in vivo* GBM model. In the mouse phantom, we were able to localize the target within 1 mm using single-projection imaging (Figure 5), even when the target was positioned as deep as 17 mm. In the *in vivo* case, fluorescence images were significantly degraded due to excitation light at 730 nm leaking into the fluorescence window, along with the presence of tissue auto-fluorescence (Figure 6a1 and b1). However, the Born ratio method effectively suppressed both the excitation light leakage and auto-fluorescence (Figure 6 a3 and b3), enabling localization of EGFR-overexpressing tumor cells to within approximately 1 mm *in vivo* (Figure 6 d-e). It is worth noting that we did not observe a significant difference in target localization between reconstructions with and without the Born ratio method in the phantom study (Section 3.2). In contrast to the *in vivo* brain imaging (Figure 6), the excitation laser at 633 nm used in the QD800 phantom study was efficiently attenuated by the mouse phantom and the 800 nm emission filter. This attenuation minimized excitation light contamination and resulted in negligible differences between the two reconstruction methods. These findings suggest that the Born ratio method is particularly beneficial in scenarios where excitation light and auto-fluorescence cannot be effectively suppressed by filters alone. Furthermore, a previous study^17^ demonstrated that the Born ratio approach not only enables robust 3D target localization but also enhances the quantification capability of FT, even in the presence of substantial background optical heterogeneity. The investigation into our FT system’s ability to quantify fluorophore concentration is currently ongoing.

As for radiation guidance, an empirical threshold of 0.5 of the maximum reconstructed value has been suggested for delineating targets in optical tomographic reconstructions. Additionally, a radiation margin has been applied to account for uncertainties in target localization and treatment delivery error for optical tomography-guided irradiation^23,27^. Based on our FT localization accuracy of 1.1 mm *in vivo* and the treatment delivery error of 0.2 mm on the SARRP^28^, we used the root sum-squared method to estimate an appropriate radiation margin. A margin of 1.2 mm is therefore recommended to be added to the FT-reconstructed target for radiation guidance with our system.

Moreover, the flexibility in fluorophore selection enhances the versatility of our FT system. A wide range of fluorescent probes^29^ can be used *in vivo*, allowing researchers to target diverse molecular markers, biological pathways, or cellular activities. This adaptability broadens the applicability of FT-guided radiation research across various cancer models and biological investigations. We anticipate that continued development of the FT-guided system will significantly enhance researchers’ ability to advance toward functional imaging-guided radiation research.

## 5. CONCLUSIONS

We developed a standalone fluorescence tomography system integrated with a small animal irradiator to enable accurate, functional imaging-guided radiation research. The system demonstrated sub-millimeter localization accuracy in both phantom and *in vivo* GBM models, with the Born ratio method effectively mitigating excitation light leakage and autofluorescence. Its modular design, compatibility with existing commercial optical system, and flexibility in fluorophore selection make it a versatile tool for diverse radiobiological studies. This work lays the foundation for advancing precision radiation targeting and functional assessment in pre-clinical radiation oncology studies.

## ACKNOWLEDGMENTS

This work was supported by the grants from National Cancer Institute, National Institutes of Health of USA (R21CA223403, R37CA230341, and R01CA240811) and Cancer Prevention and Research Institute of Texas RR200042, and RP180770. We thank Ciara Newman for helping with cell culture for the *in vivo* experiment.

